# Adaptive mutations at lysine residues of PRRSV nsp12 enable evasion of host proteasomal and selective autophagic degradation

**DOI:** 10.1101/2025.09.02.673672

**Authors:** Yongjie Chen, Zhan He, Zishen Chen, Ling Huang, Haotong Lu, Siyong Zeng, Baoying Huang, Chunhe Guo

## Abstract

Ubiquitin signaling in viral infections can either enhance immune responses and degrade viral proteins or be exploited by viruses to target host antiviral factors. Porcine reproductive and respiratory syndrome virus (PRRSV) poses a significant threat to the pig industry, with its non-structural proteins (nsps) critical for virulence and replication. Here, we investigated the role of ubiquitination in the stability of PRRSV-encoded nsps and found that nsp12 was specifically degraded via the ubiquitin-proteasome system. Mechanistically, nsp12 underwent K48- and K63-linked polyubiquitination at lysine residues 89, 91, 127, and 130. RNF114 served as an E3 ubiquitin ligase for nsp12, with its enzymatic activity essential for both nsp12 degradation and viral replication. Additionally, nsp12 underwent ubiquitin-dependent selective autophagy through receptor-mediated recognition, wherein NBR1, SQSTM1, and NDP52 bridged its interaction with LC3 for autophagic degradation. Evolutionary analyses revealed that PRRSV nsp12 acquired non-lysine residues at positions 89, 127, and 130 during viral adaptation. Correspondingly, recombinant PRRSV strains carrying the K91/127/130R mutations within nsp12 exhibited enhanced replication, while a revertant strain with the R89K mutation in nsp12 showed attenuated infectivity. Mass spectrometry analysis further identified significant enrichment of ubiquitination-related modifications among nsp12-interacting proteins. These findings provide valuable insights for anti-PRRSV drug design and highlight the challenge posed by adaptive mutations in viral proteins to the swine industry.

**Author Summary:** Ubiquitination plays crucial roles in both proteasomal degradation and selective autophagy during viral infections, yet its impact on the stability of porcine reproductive and respiratory syndrome virus (PRRSV) nonstructural proteins (nsps) remains unexplored. Here, we found that nsp12 was targeted for proteasomal degradation through K48/K63-linked polyubiquitination at lysine residues 89, 91, 127, and 130, which requires the involvement of E3 ubiquitin ligase RNF114. Furthermore, nsp12 underwent receptor-mediated selective autophagic degradation through the action of NBR1, SQSTM1, and NDP52. Using reverse genetics technology, mutations of lysine to arginine in PRRSV nsp12 enhanced viral replication, whereas the reverse mutations reduced its infectivity. Our findings demonstrate that PRRSV evades host degradation by acquiring adaptive mutations within nsp12, which counteract ubiquitin-dependent clearance thus enhancing viral fitness. This novel mechanism illustrates a key viral immune evasion strategy and underlines the challenge in controlling PRRSV.

## Introduction

Livestock farming is a crucial pillar of the global food supply chain, yet bacterial and viral diseases pose significant challenges to its development [1]. These diseases not only lead to animal mortality and reduced production performance, resulting in substantial economic losses, but also pose potential threats to food safety [2,3]. Since its first report in the United States in 1987, porcine reproductive and respiratory syndrome (PRRS) has spread worldwide, including China, severely impeding the development of the pig farming industry and causing significant economic losses [4,5]. PRRS virus (PRRSV) is an enveloped single-stranded positive-sense RNA virus that belongs to the family *Arteriviridae* and the genus *Betaarterivirus* [6,7]. The genome of PRRSV is approximately 15 kb in length and contains at least 11 open reading frames (ORFs), encoding at least 16 non-structural proteins (nsps) and 8 structural proteins [8,9]. Pigs of all ages are susceptible to PRRSV, with pregnant sows and newborn piglets being the most susceptible [10]. PRRSV primarily infects macrophages in pigs, particularly porcine alveolar macrophages (PAMs). After replicating within macrophages, the virus disseminates to lymphatic tissues and the lungs [11]. It causes severe reproductive disorders in sows, respiratory diseases in piglets, and secondary infections [12]. As an RNA virus, PRRSV evolves rapidly due to its high mutation rate and recombination frequency, and is capable of rapidly generating new mutant strains [13]. Genomic variation allows PRRSV to easily evade host defense and recognition by the immune system, posing significant challenges for epidemic prevention and control [14].

In recent years, with the growing threat posed by emerging pathogens to public health security and the lack of effective drugs and vaccines for certain traditional pathogens, finding new therapeutic targets has become a global research focus [15,16]. In this context, ubiquitination modification has gradually gained widespread attention. Ubiquitination, a crucial form of posttranslational modification, plays a vital role in protein activity, protein-protein interactions, and subcellular localization of proteins [17]. This modification not only influences the function of individual proteins but also extensively participates in the regulation of nearly all life activities, including the cell cycle, proliferation, apoptosis, differentiation, signal transduction, injury repair, inflammatory immunity, and more [18–20]. In recent years, with the deepening of research on ubiquitination, its functions in host immunity and viral infections have gradually garnered attention. Ubiquitination is involved in both innate and adaptive immune responses by regulating the activation, proliferation, and differentiation, as well as the production and release of immune molecules [20,21]. Viruses can exploit the host’s ubiquitination system to evade immune surveillance, modulate the stability and activity of viral proteins, as well as the assembly and release of viral particles, thereby promoting viral replication and dissemination [22–24]. Conversely, to counteract viral infections, the host utilizes ubiquitination to modify viral proteins, thereby targeting and degrading key viral proteins or host cell proteins hijacked by the virus, ultimately influencing viral infection and replication [25,26]. When the proteasome system is overloaded, ubiquitinated proteins are degraded through the selective autophagy pathway [27]. In selective autophagy, ubiquitination serves as the core mechanism for recognition and degradation of specific substrates. Upon being ubiquitinated, proteins are recognized by specific selective autophagy receptors. These receptors, such as NBR1(NBR1 autophagy cargo receptor), OPTN (optineurin), SQSTM1/p62 (sequestosome 1), CALCOCO2/NDP52 (calcium binding and coiled- coil domain 2) and TOLLIP (toll interacting protein), play a bridging role in the autophagy process [28]. They not only recognize ubiquitinated proteins but also bind to LC3 (light chain 3) during autophagosome formation, to ensure that the proteins are correctly directed to the autophagosomes for degradation. Ultimately, these proteins are degraded through the autophagosome-lysosome pathway [29].

In this study, we found that the nsp12 of PRRSV was unstable in host cells. It underwent polyubiquitination and was subsequently degraded via both the ubiquitin-proteasome pathway and selective autophagy. To evade these host defense mechanisms, PRRSV had acquired adaptive mutations at lysine residues within nsp12 during genetic evolution, thereby enhancing its resistance to host degradation and facilitating more efficient virus replication. Our research not only reveals the crucial roles of the ubiquitin-proteasome system and selective autophagy in host antiviral defense but also elucidates how viruses escape these defenses through adaptive evolution.

## Results

### Nsp12 undergoes degradation via the ubiquitin-proteasome pathway

To investigate whether the stability of PRRSV-encoded nsps is regulated by the ubiquitin-proteasome pathway, we examined the effect of the proteasome inhibitor MG132 on the abundance of viral nsps. We found that MG132 treatment specifically and significantly elevated the protein levels of nsp12, but did not affect other nsps (Figure 1A), as confirmed by fluorescence intensity measurements (Figure S1A). Additionally, nsp12 expression increased in a concentration-dependent manner upon MG132 treatment (Figures 1B and S1B). To further investigate nsp12 stability, we used cycloheximide (CHX), an inhibitor of eukaryotic translation elongation and protein synthesis. CHX treatment resulted in the rapid degradation of nsp12, while MG132 treatment significantly stabilized its protein levels (Figure 1C). These results demonstrate that nsp12 is unstable and can be degraded via the proteasome pathway. For a protein to be recognized and degraded by the proteasome, it must first undergo ubiquitination. Ubiquitination is a prerequisite for proteasomal degradation and a core mechanism for regulating protein homeostasis within cells. To investigate whether nsp12 undergoes ubiquitination, we performed co-immunoprecipitation (Co-IP) assays. The results showed that overexpressed ubiquitin bound to the nsp12, forming polyubiquitin chains (Figure 1D). Overexpression of nsp12 also induced ubiquitin translocation from the nucleus to the cytoplasm, where it colocalized with nsp12 (Figure 1E). Additionally, endogenous ubiquitin was found to bind to nsp12 (Figure 1F), further confirming that nsp12 is ubiquitinated within cells. To determine the types of ubiquitination on nsp12, we coexpressed nsp12-mCherry with hemagglutinin (HA)- ubiquitin wild-type (HA-Ub-WT) or lysine-specific mutants (K6, K11, K27, K29, K33, K48, K63, K48R and K63R). The results demonstrated that nsp12 undergoes K48- and K63-linked polyubiquitination (Figure 1G and H). In contrast, cotransfected with the HA-Ub-K48R and HA-Ub-K63R plasmids significantly reduced the ubiquitination of nsp12 (Figure 1I). These findings confirm that K48- and K63-linked polyubiquitination mediates the recognition and subsequent degradation of nsp12 by the proteasome.

**Figure 1.**
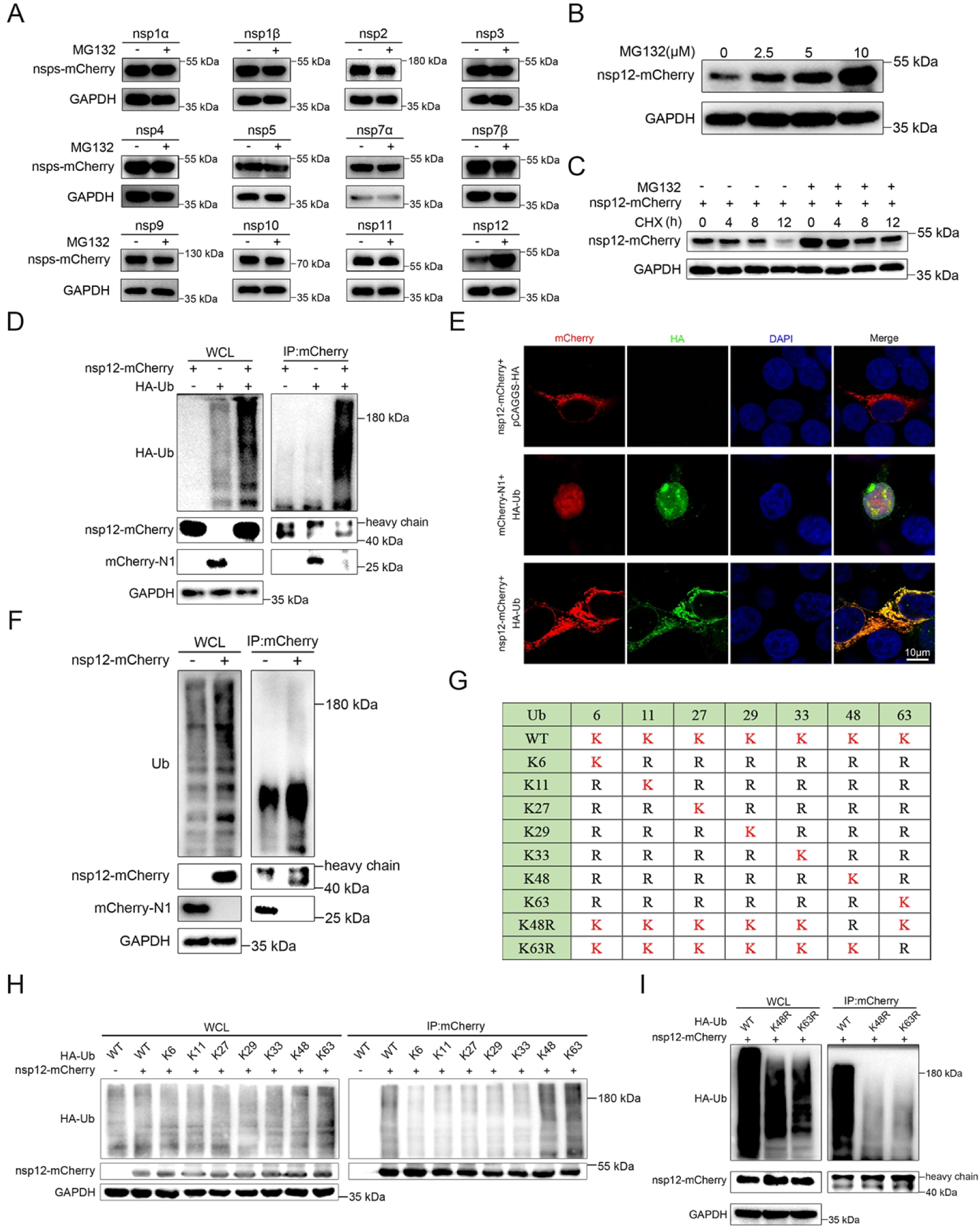
Nsp12 undergoes degradation via the ubiquitin-proteasome pathway. (**A**) HEK293T cells were transfected with PRRSV nsps-mCherry plasmids and treated with MG132 (10 μM) or DMSO for 12 h prior to harvest. Protein levels of nsps were detected by western blotting. (**B**) HEK293T cells were transfected with nsp12-mCherry and treated with different concentrations of MG132 for 12 h before harvesting. Protein levels of nsp12 were analyzed by western blotting. (**C**) HEK293T cells were transfected with the nsp12-mCherry for 12 h, then treated with or without MG132 (10 μM) for 12 h, followed by the addition of CHX (50 μg/mL). Cells were harvested at different time points, and the cell lysates were analyzed. (**D and E**) HEK293T cells were transfected with nsp12-mCherry or an empty vector, along with HA-Ub or an empty vector. Co-IP was performed using anti-mCherry beads, followed by western blotting (**D**). Confocal assays were used to assess the colocalization of nsp12 and Ub (**E**). (**F**) HEK293T cells were transfected with nsp12-mCherry alone. Whole cell lysates (WCL) were incubated with anti-mCherry beads and subjected to western blotting. (**G**) Schematic diagram of Ub mutants (K6, K11, K27, K29, K33, K48, K63, K48R and K63R). (**H and I**) HEK293T cells were transfected with nsp12 and Ub-WT or Ub mutants. Co-IP was performed using anti-mCherry beads, followed by western blotting.

### Ubiquitin-proteasome degradation of PRRSV nsp12 via K89/K91/K127/K130 ubiquitination

To further identify the ubiquitination sites on nsp12, we analyzed its full-length sequence and identified seven lysine residues. These residues were individually mutated to arginine to assess their role in ubiquitination (Figure 2A). The results showed that treatment with MG132 significantly increased the expression levels of all nsp12 mutants, suggesting that multiple lysine residues are crucial for the ubiquitination and degradation of nsp12 (Figure 2B and C). We then generated the K0 mutant by mutating all seven lysine residues to arginine. Unlike nsp12-WT, MG132 treatment did not increase the protein levels of the nsp12-K0 mutant (Figure 2D). When cells overexpressing nsp12-WT and nsp12-K0 were treated with CHX, nsp12-K0 exhibited significantly higher protein stability than nsp12-WT (Figure 2E). Moreover, ubiquitination levels of nsp12-K0 were nearly undetectable compared to nsp12-WT (Figure 2F). To identify specific ubiquitination sites, we individually restored lysine in the K0 mutant backbone (Figure 2G). The results indicated that the protein levels of nsp12^K89^, nsp12^K91^, nsp12^K127^, and nsp12^K130^ mutants were significantly reduced, but were stabilized upon MG132 treatment (Figure 2H and I). Notably, when lysines at positions 127 and 130 were simultaneously mutated to arginine (nsp12^K127/130R^), MG132 treatment still increased the protein levels. However, when all four lysines (K89, K91, K127, and K130) were mutated to arginine (nsp12^K89/91/127/130R^), MG132 treatment failed to increase the protein levels (Figure 2J). CHX treatment further revealed that nsp12^K89/91/127/130R^ displayed significantly higher protein stability than nsp12-WT (Figure 2K). Additionally, nsp12^K89/91/127/130R^ exhibited significantly fewer polyubiquitin chains than nsp12-WT (Figure 2L). In summary, these results demonstrate that nsp12 undergoes ubiquitin-proteasomal degradation mediated by ubiquitination at lysine residues 89, 91, 127, and 130.

**Figure 2.**
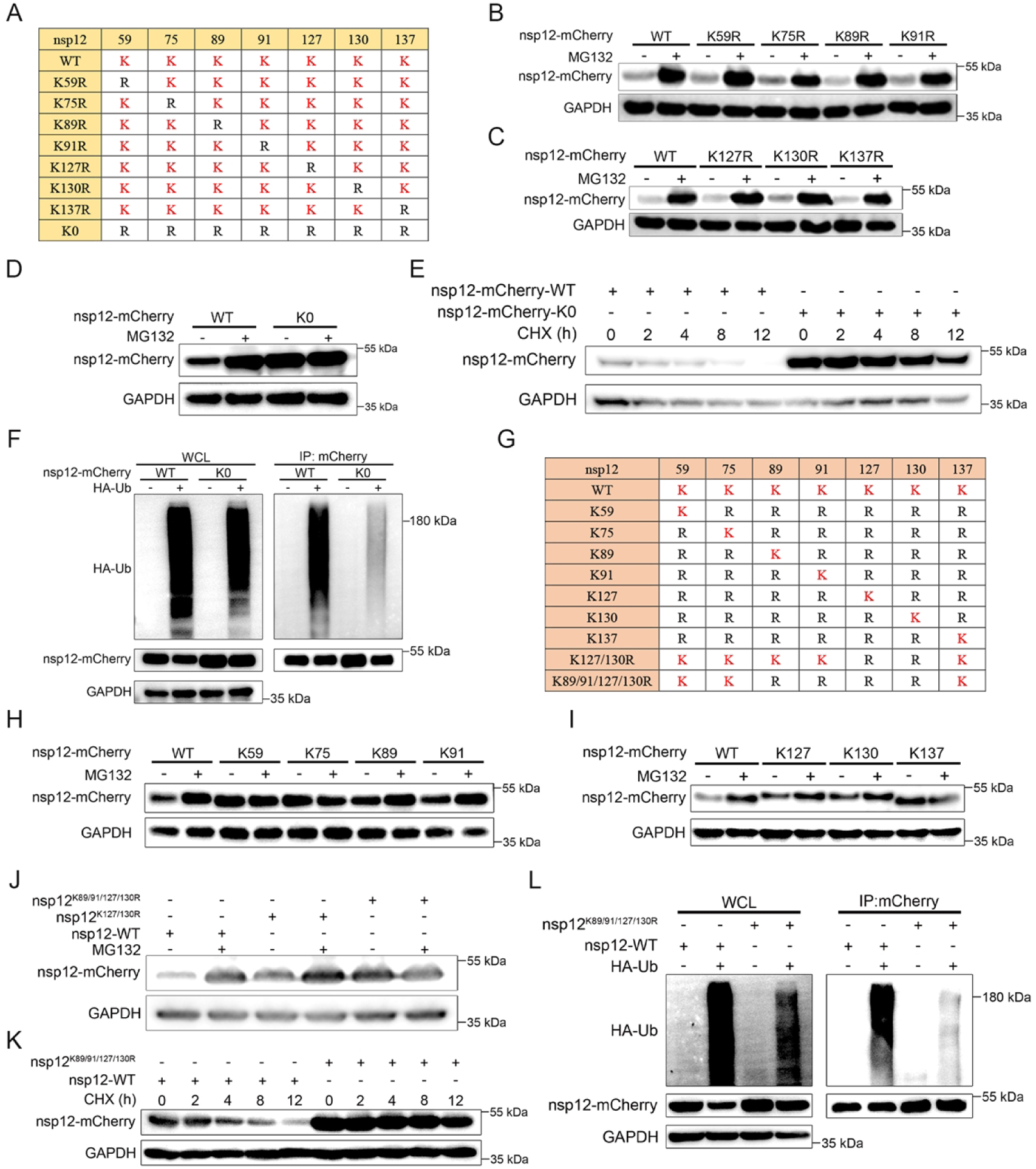
**Ubiquitin-proteasome degradation of PRRSV nsp12 via K89/K91/K127/K130 ubiquitination**. (**A**) Schematic diagram of the construction strategy for nsp12 lysine-to-arginine mutations. (**B and C**) HEK293T cells were transfected with nsp12 or nsp12 carrying lysine mutants and treated with or without MG132 (10 μM) for 12 h prior to harvest. Protein levels of nsp12 were detected by western blotting. (**D**) HEK293T cells were transfected with nsp12-WT or nsp12-K0. Cells were treated with or without MG132 (10 μM) for 12 h prior to harvest. Protein levels of nsp12 were detected by western blotting. (**E**) HEK293T cells were transfected with nsp12-WT or nsp12-K0 for 24 h, followed by the addition of CHX (50 μg/mL). Cells were harvested at different time points, and the cell lysates were analyzed. (**F**) HEK293T cells were transfected with nsp12-WT or nsp12-K0, along with HA-Ub or an empty vector. Co-IP was performed using anti-mCherry beads, followed by western blotting. (**G**) Schematic diagram illustrating the use of the K0 mutant as a backbone, restoring arginine to lysine. (**H and I**) HEK293T cells were transfected with nsp12-WT or nsp12 single mutants where arginine was replaced by lysine based on the K0 mutant. Cells were treated with or without MG132 (10 μM) for 12 h prior to harvest. Protein levels of nsp12 were detected by western blotting. (**J**) HEK293T cells were transfected with plasmids expressing nsp12^K127/130R^ or nsp12^K89/91/127/130R^. Cells were treated with or without MG132 (10 μM) for 12 h prior to harvest. Protein levels of nsp12 were detected by western blotting. (**K**) HEK293T cells were transfected with nsp12-WT or nsp12^K89/91/127/130R^ for 24 h, followed by the addition of CHX (50 μg/mL). Cells were harvested at different time points, and cell lysates were analyzed. (**L**) HEK293T cells were transfected with nsp12-WT or nsp12^K89/91/127/130R^, along with HA-Ub or an empty vector. Co-IP was performed using anti-mCherry beads, followed by western blotting.

### RNF114 specifically ubiquitinates nsp12 at K127 and K130 in a manner dependent on its E3 ubiquitin ligase activity

Previous studies have shown that RNF114, an E3 ubiquitin ligase, interacts with nsp12 and promotes its degradation via the proteasomal pathway [30]. However, the precise underlying mechanism remains unclear. In this study, we similarly observed that nsp12 interacted with RNF114 and underwent RNF114-mediated degradation through the ubiquitin-proteasome pathway (Figure 3A-C). The genetic diversity of PRRSV strains presents a significant challenge for effective control. To assess whether RNF114- mediated degradation of nsp12 could broadly inhibit PRRSV replication across different strains, we transfected cells with increasing doses of RNF114 and subsequently infected them with various PRRSV strains (CH-1a, TA-12, JXA1, SD16, and WUH3). Our results showed a dose-dependent reduction in both viral N protein and nsp1α protein levels following RNF114 overexpression (Figure S2A-E). Consistent with these findings, fluorescence intensity measurements revealed a significant decrease in PRRSV N protein signals, corresponding to higher RNF114 expression levels (Figure S2F-J). These results collectively suggest that RNF114 exhibits broad-spectrum antiviral activity against multiple PRRSV strains.

**Figure 3.**
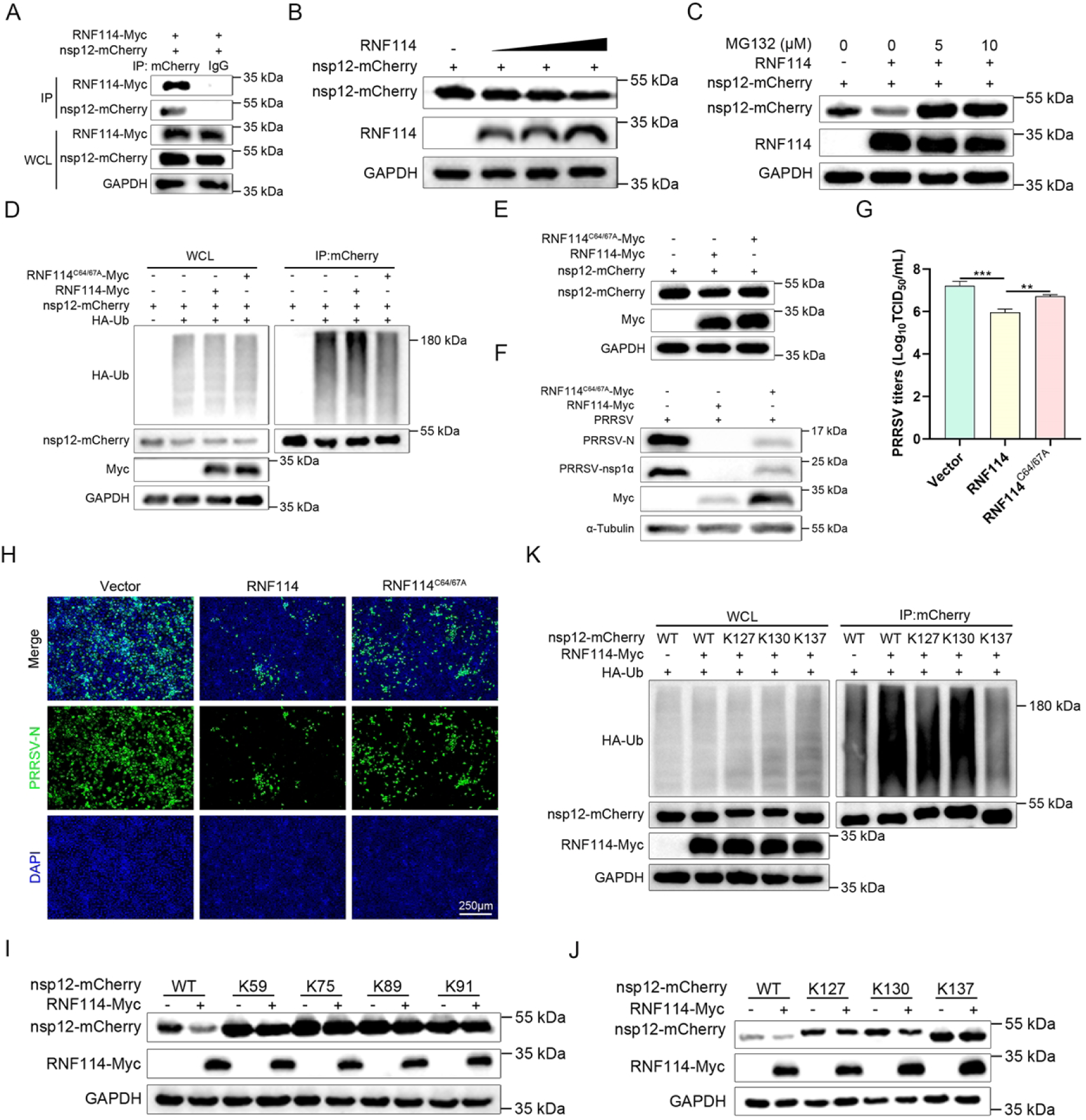
RNF114 specifically ubiquitinates nsp12 at K127 and K130 in a manner dependent on its E3 ubiquitin ligase activity. (**A**) HEK293T cells were cotransfected with nsp12 and RNF114. Co-IP was performed using anti-mCherry or anti-IgG beads, followed by western blotting. (**B**) HEK293 cells were cotransfected with nsp12 and increasing amounts of RNF114, followed by western blotting. (**C**) HEK293T cells expressing nsp12 with or without RNF114 were treated with indicated concentrations of MG132 for 12 h before harvesting for western blotting. (**D**) HEK293T cells were cotransfected with nsp12 and either RNF114 or RNF114^C64/67A^, with or without HA- Ub. Co-IP was performed using anti-mCherry beads. (**E**) HEK293T cells were cotransfected with nsp12 and either RNF114 or RNF114^C64/67A^, followed by western blotting. (**F-H**) Marc-145 cells were transfected with RNF114 or RNF114^C64/67A^ for 24 h, then infected with PRRSV strain CH-1a at a multiplicity of infection (MOI) of 0.5 for 24 h. The viral N and nsp1α protein levels were measured by western blotting (**F**). Viral titers in supernatants were quantified by TCID_50_ (**G**). N protein expression was assessed by IFA (**H**). (**I and J**) HEK293T cells were transfected with nsp12-WT or single-lysine nsp12 mutants, with or without RNF114, followed by western blotting. (**K**) HEK293T cells were cotransfected with nsp12-WT or nsp12 mutants (nsp12^K127^, nsp12^K130^ or nsp12^K137^) along with HA-Ub and either RNF114 or empty vector. Co-IP was performed using anti-mCherry beads.

To determine whether RNF114’s regulatory effect on nsp12 depends on its E3 ubiquitin ligase activity, we investigated the role of its catalytic residues. It has been established that cysteine residues at positions 64 and 67 (C64 and C67) are critical for RNF114’s ligase activity [31]. Our results showed that mutation of these residues abolished RNF114’s ability to ubiquitinate nsp12 (Figure 3D), impairing nsp12 degradation (Figure 3E). Furthermore, the catalytically inactive RNF114 mutant exhibited a significantly reduced capacity to antagonize PRRSV replication (Figure 3F-H). To determine the specific ubiquitination sites on nsp12 mediated by RNF114, we performed mutagenesis on seven lysine residues of nsp12. We found that only nsp12^K127^ or nsp12^K130^ remained susceptible to RNF114-induced degradation, whereas all other mutants exhibited resistance (Figure 3I and J). Additionally, we demonstrated that RNF114 enhanced ubiquitination of nsp12 at K127 and K130, but not at K137 (Figure 3K). These findings indicate that K127 and K130 are the key ubiquitination sites on nsp12 targeted by RNF114. In summary, our study demonstrates that RNF114, via its E3 ubiquitin ligase activity, specifically ubiquitinates nsp12 at K127 and K130, thereby promoting its proteasomal degradation.

### Ubiquitin-dependent selective autophagy mediates the degradation of nsp12 in host cells

So far, we have demonstrated that host cells degrade nsp12 through the proteasome pathway, as evidenced by the stabilization of nsp12 upon MG132 treatment. To further investigate whether exogenous ubiquitin affects nsp12 expression, human embryonic kidney 293T (HEK293T) cells were transfected with nsp12 in the presence or absence of HA-Ub. Exogenous ubiquitin promoted nsp12 degradation in a dose-dependent manner (Figure 4A and B), whereas nsp12-K0 mutant remained unaffected (Figure 4C), confirming the requirement of ubiquitination for this process. It is noteworthy that ubiquitin-mediated nsp12 degradation was suppressed not only by the proteasome inhibitor MG132 but also by the autophagy inhibitor 3-MA (Figure 4D). This was further supported by the pronounced co-localization of nsp12-mCherry, HA-Ub, and GFP-LC3 (Figure 4E). The involvement of autophagy was validated through dose-dependent stabilization of nsp12 by 3-MA (Figure 4F) and CHX chase assays, which revealed a significantly extended nsp12 half-life upon 3-MA treatment (Figure 4G). To identify the lysine residues critical for ubiquitin-dependent autophagic degradation, we treated a series of nsp12 lysine mutants (each containing a single lysine-to-arginine substitution) with 3-MA. While 3-MA restored protein levels for all nsp12 mutants (Figure 4H and I), it had no effect on the nsp12-K0 mutant (Figure 4J), indicating that ubiquitination at one or more lysine residues is essential for autophagic targeting. Further analysis of nsp12 mutants retaining individual lysines revealed that residues K89, K91, K127, and K130 were each sufficient to mediate ubiquitin-dependent autophagic degradation, as their protein levels were significantly rescued by 3-MA (Figure 4K and L). Collectively, these results demonstrate that nsp12 is regulated by both the ubiquitin-proteasome system and ubiquitin-dependent selective autophagy, with specific lysine residues (K89, K91, K127, and K130) serving as critical determinants of its turnover.

**Figure 4.**
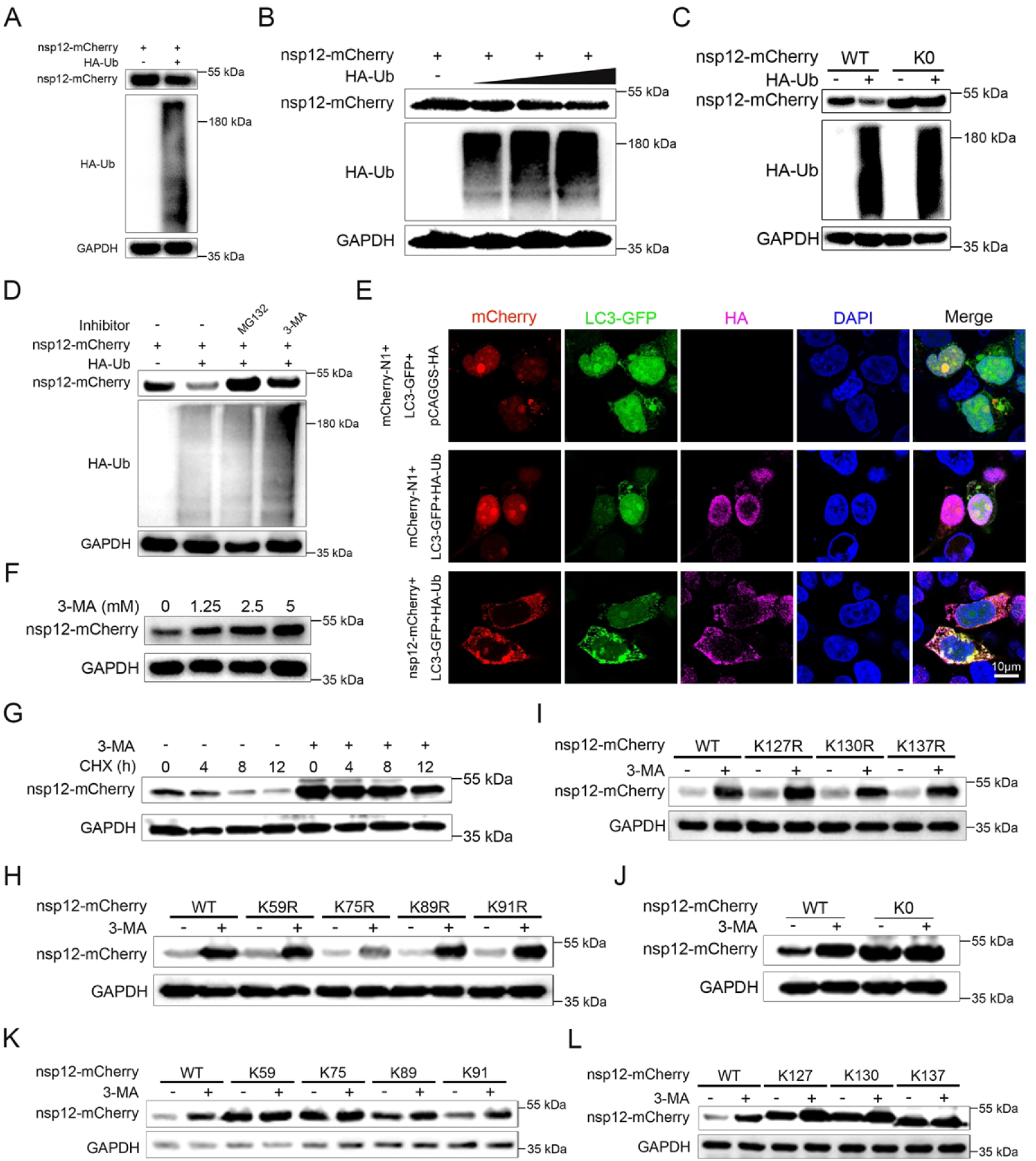
Ubiquitin-dependent selective autophagy mediates nsp12 degradation in host cells. (**A**) HEK293 cells were cotransfected with nsp12 and HA-Ub, followed by western blotting. (**B**) HEK293 cells were cotransfected with nsp12 and different amounts of HA-Ub, followed by western blotting. (**C**) HEK293T cells were transfected with nsp12-WT or nsp12-K0, along with HA-Ub or an empty vector. Cell samples were harvested for western blotting. (**D**) HEK293T cells transfected with nsp12 and HA-Ub were treated with MG132 (10 μM) or 3-MA (10 mM) for 12 h prior to harvest, followed by western blotting. (**E**) HEK293T cells were transfected with nsp12-mCherry or an empty vector, along with HA-Ub or an empty vector, and GFP-LC3. Confocal microscopy was used to assess the colocalization of nsp12, Ub, and LC3. (**F**) HEK293T cells were transfected with nsp12-mCherry plasmids and treated with different concentrations of 3-MA for 12 h prior to harvest, followed by western blotting. (**G**) HEK293T cells transfected with nsp12-mCherry for 12 h were treated with or without 3-MA (10 mM) for 12 h, followed by the addition of CHX (50 μg/mL). Cells were harvested at different time points, and the cell lysates were analyzed. (**H and I**) HEK293T cells transfected with nsp12 or nsp12 carrying lysine mutants were treated with or without 3-MA (5 mM) for 12 h prior to harvest. Protein levels of nsp12 were detected using western blotting. (**J**) HEK293T cells were transfected with nsp12-WT or nsp12-K0. Cells were treated with or without 3-MA (10 mM) for 12 h prior to harvest. Cells were harvested for western blotting. (**K and L**) HEK293T cells were transfected with nsp12-WT or single-lysine nsp12 mutants. Cells were treated with or without 3- MA (5 mM) for 12 h prior to harvest. Protein levels of nsp12 were detected by western blotting.

### Nsp12 interacts with multiple autophagy receptors

In selective autophagy, ubiquitination serves as the central recognition signal for substrate degradation. Ubiquitinated proteins are typically recognized by selective autophagy receptors, including NBR1, SQSTM1, NDP52, and others, which simultaneously bind LC3 to facilitate autophagosome formation. This dual recognition mechanism ensures precise targeting of substrates for autophagic degradation. Next, we investigated whether nsp12 interacts with autophagy receptors. Co-IP assays demonstrated that nsp12 associates with multiple autophagy receptors, including NBR1, OPTN, SQSTM1, NDP52, and TOLLIP (Figure 5A). Confocal microscopy further revealed significant co-localization between nsp12 and these receptors (Figure 5B). Interestingly, coexpression with NBR1, OPTN, or TOLLIP induced a striking redistribution of nsp12 from a diffuse cytoplasmic pattern to distinct aggregates, suggesting these receptors may regulate nsp12’s subcellular localization and function. To distinguish between direct and indirect interactions, we performed yeast two-hybrid assays, which confirmed direct binding between nsp12 and NBR1, NDP52, and TOLLIP (Figure 5C and D). These physical interactions were further supported by computational modeling (Figure S3B-D). These findings suggest that nsp12 may undergo selective autophagic degradation through both direct and indirect interactions with multiple autophagy receptors.

**Figure 5.**
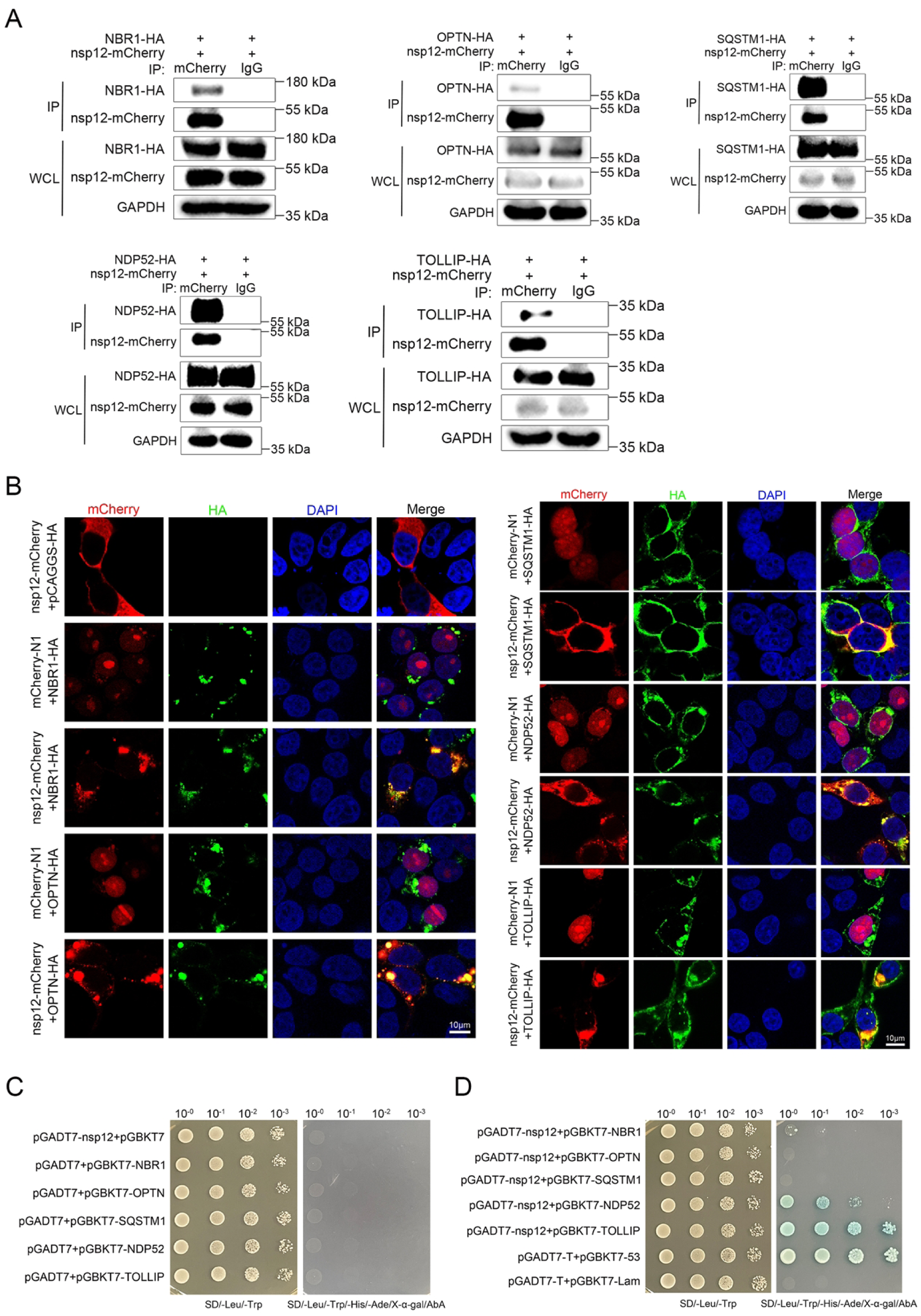
Nsp12 interacts with multiple autophagy receptors. (**A**) HEK293T cells were cotransfected with nsp12 and NBR1, OPTN, SQSTM1, NDP52, or TOLLIP. Co- IP was performed using anti-mCherry or anti-IgG beads, followed by western blotting. (**B**) HEK293T cells were transfected with NBR1, OPTN, SQSTM1, NDP52, and TOLLIP, along with nsp12 or an empty vector. Confocal microscopy was used to assess their colocalization. (**C and D**) The bait and prey plasmids were cotransfected into yeast cells to detect potential interactions. Positive control group: pGBKT7-53 and pGADT7-T. Negative control group: pGBKT7-Lam and pGADT7-T. SD/-Leu/-Trp/-His/-Ade, synthetic dropout (SD) agar medium without leucine, tryptophan, histidine, and adenine; SD/-Leu/-Trp, SD agar medium without leucine and tryptophan.

### Nsp12 is degraded through the selective autophagy pathway mediated by NBR1, SQSTM1, and NDP52

As autophagy receptors typically bridge connecting ubiquitinated substrates to LC3, we first confirmed that nsp12 formed complexes with both LC3 and multiple autophagy receptors (NBR1, OPTN, SQSTM1, NDP52, and TOLLIP) through Co-IP assays (Figure 6A). Immunofluorescence assays further demonstrated robust co-localization of these three components (Figure 6B). While Co-IP and confocal microscopy confirmed the nsp12-LC3 interaction (Figure 6C and D), yeast two-hybrid assays revealed no direct binding (Figure 6E), suggesting this interaction requires autophagy receptors as molecular adaptors. To identify the specific receptors responsible for nsp12 degradation, we performed targeted knockdown experiments. Silencing of NBR1, SQSTM1, or NDP52 significantly restored nsp12 protein levels, whereas OPTN and TOLLIP depletion showed no effect (Figure 6F). More importantly, under ubiquitin- overexpressing conditions, only the knockdown of NBR1, SQSTM1, or NDP52 abolished ubiquitin-induced nsp12 degradation (Figure 6G and H). These results demonstrate that while nsp12 interacts with multiple autophagy receptors, its selective autophagic degradation specifically depends on NBR1, SQSTM1, and NDP52.

**Figure 6.**
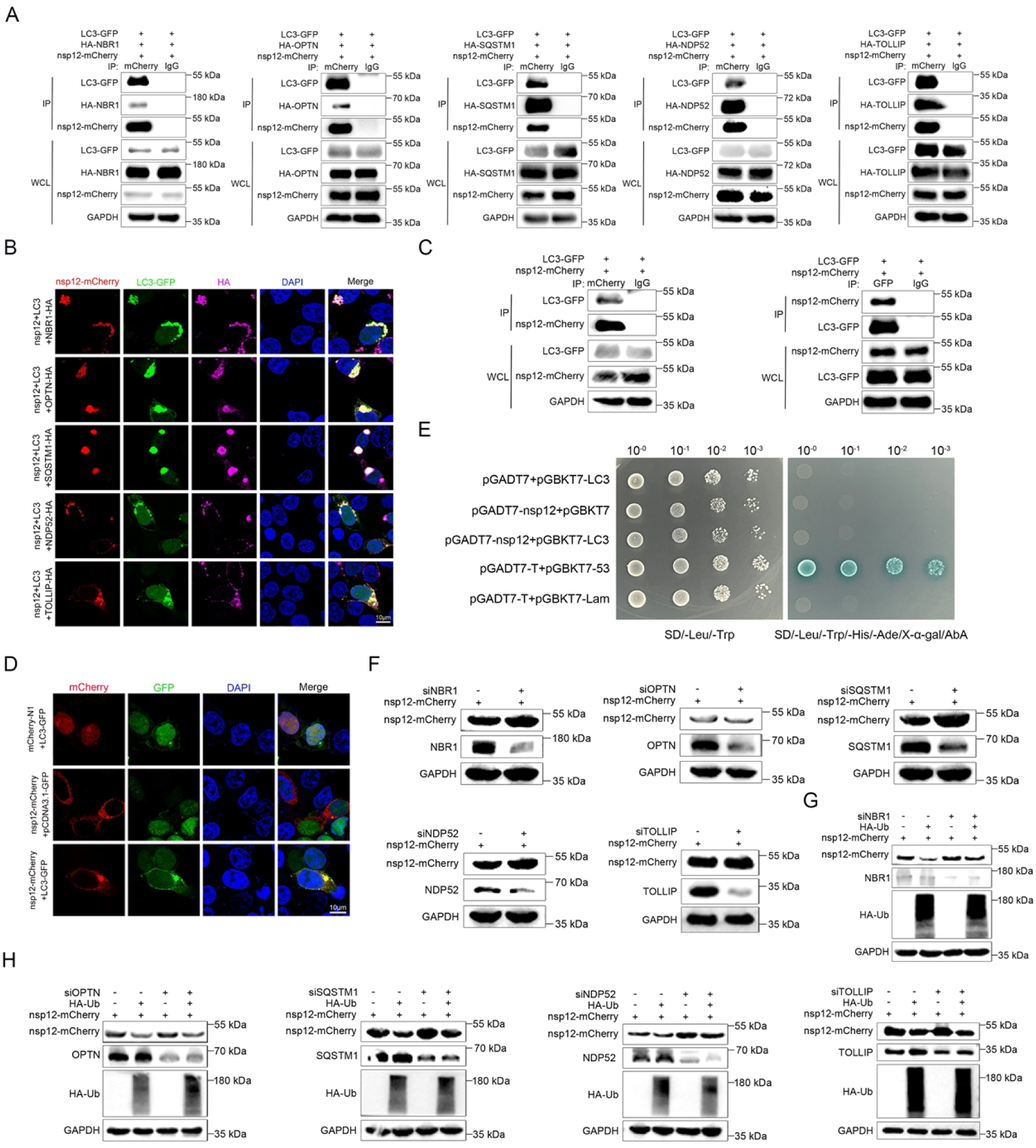
Nsp12 is degraded through the selective autophagy pathway mediated by NBR1, SQSTM1, and NDP52. (**A and B**) HEK293T cells were cotransfected with plasmids encoding nsp12-mCherry, LC3-GFP, and HA-tagged autophagy receptors (NBR1, OPTN, SQSTM1, NDP52, and TOLLIP). Co-IP was performed using anti- mCherry or anti-IgG beads, followed by western blotting analysis (**A**). Colocalization was assessed by confocal microscopy (**B**). (**C and D**) HEK293T cells were cotransfected with plasmids encoding nsp12-mCherry and LC3-GFP. Co-IP was performed using anti-mCherry, anti-GFP or anti-IgG beads, followed by western blotting (**C**). Confocal assays were used to assess the colocalization between nsp12 and LC3 (**D**). (**E**) The pGADT7-nsp12 and pGBKT7-LC3 plasmids were cotransfected into yeast cells to assess protein-protein interactions. (**F**) HEK293T cells were transfected with siRNAs targeting NBR1, OPTN, SQSTM1, NDP52, or TOLLIP for 12 h, followed by transfection with nsp12-mCherry for 24 h. Scrambled siRNAs were included as controls. Cell lysates were analyzed by western blotting. (**G and H**) HEK293T cells were transfected with siRNAs targeting NBR1, OPTN, SQSTM1, NDP52, or TOLLIP for 12 h, followed by transfection with nsp12-mCherry, along with HA-Ub or an empty vector for 24 h. Scrambled siRNAs were included as controls. Cell lysates were analyzed by western blotting.

### Multiple lysine residues within nsp12 show a tendency to mutate to arginine during the genetic evolution of PRRSV

To adapt to environmental pressures and evade host defenses, viral proteins often acquire adaptive mutations at key amino acid sites that enhance their survival and replication. To investigate the genetic evolution of PRRSV nsp12, we performed phylogenetic analysis using nsp12 sequences from the GenBank database (Table S1). The results revealed that Lineages 1 and 8 are the predominant circulating strains of PRRSV-2 (Figure 7A). Notably, most strains submitted between 2020 to 2024 clustered within these Lineages, further confirming their dominance (Figure 7A). Comparative nucleotide and amino acid homology analyses of nsp12 across four lineages demonstrated substantial genetic variability. Lineage 1 exhibited the highest diversity (nucleotide homology: 81.7%∼100.0%; amino acid homology: 87.6%∼100.0%), while Lineage 5 showed the most conservation (nucleotide homology: 99.3%∼99.8%; amino acid homology: 99.3%∼100.0%) (Figure 7B). Notably, we identified a recurring pattern of lysine-to-arginine at positions 59, 89, 127, 130, and 137 in circulating strains relative to the ancestral VR2332 strain (Figure 7C and D, Table S2). Strikingly, three of lysine residues (89, 127, and 130) correspond to experimentally validated ubiquitination sites. This observation suggests that PRRSV may escape host-mediated ubiquitination-dependent degradation by substituting lysine with arginine at these critical positions during evolution, thereby enhancing its resistance to host antiviral mechanisms.

**Figure 7.**
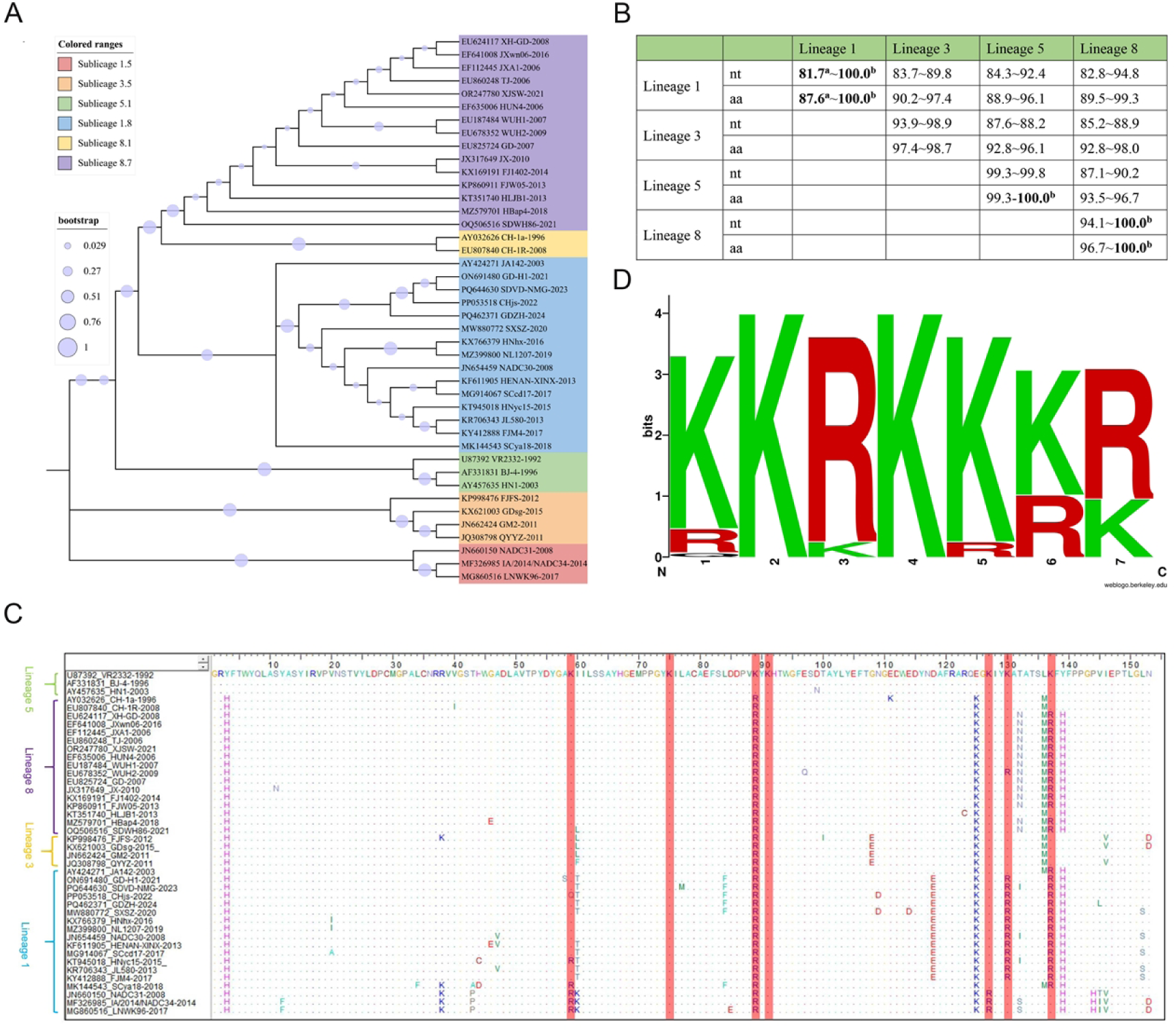
Multiple lysine residues within nsp12 show a tendency to mutate to arginine during the genetic evolution of PRRSV. (**A**) Phylogenetic analysis of nsp12. The phylogenetic tree was constructed using the NJ method in MEGA software with 1,000 bootstrap replicates. (**B**) Nucleotide and amino acid homology analysis based on the nsp12 (%). ^a^Indicates the lowest homology; ^b^Indicates the highest homology. (**C**) Alignment of nsp12 amino acid sequences from different PRRSV strains. (**D**) The frequency of lysine mutations at positions 59, 75, 89, 91, 127, 130, and 137 in PRRSV nsp12 was analyzed using the website (http://weblogo.berkeley.edu/).

### The rPRRSV-nsp12^K91/127/130R^ exhibits greater replication capacity compared to rPRRSV

To investigate whether PRRSV nsp12 evades host-mediated degradation through adaptive mutations, we compared the lysine residues of the ancestral VR2332 strain with the contemporary TA-12 strain. Notably, lysine residues at positions 91, 127, and 130 were highly conserved (Figure S4A). Since single mutations at these sites were previously shown to be insufficient for complete immune evasion, we generated a recombinant strain (rPRRSV-nsp12^K91/127/130R^) using the TA-12 backbone (Figure S4B). Following rescue in HEK293T cells, western blot analysis at 48 h post-transfection revealed significantly higher expression of PRRSV nsp1α in cells transfected with the triple mutant compared to wild-type controls (Figure 8A), suggesting that these mutations may alter viral gene expression. The successful rescue of rPRRSV- nsp12^K91/127/130R^ was confirmed through cytopathic effect (CPE) (Figure S4C), indirect immunofluorescence assay (IFA) (Figure S4D and E), western blotting (Figure S4F), reverse transcription (RT)-PCR (Figure S4G), and Sanger sequencing (Figure S4H). Subsequently, we evaluated the impact of these lysine mutations on PRRSV replication in Marc-145 cells. The recombinant rPRRSV-nsp12^K91/127/130R^ induced more pronounced CPE than rPRRSV (Figure 8B). Additionally, we observed significantly higher mRNA and protein levels of the N and nsp1α (Figure 8C-E), as well as increased fluorescence intensity for rPRRSV-nsp12^K91/127/130R^ (Figure 8F and G), compared to rPRRSV. Although both strains formed similar-sized plaques, rPRRSV- nsp12^K91/127/130R^ exhibited a higher viral titer than rPRRSV (Figure 8H and I). Moreover, infection with rPRRSV-nsp12^K91/127/130R^ resulted in more lactate dehydrogenase (LDH) release compared to rPRRSV (Figure 8J). Importantly, we observed a significant increase in the proportion of strains with lysine-to-arginine mutations at positions 127 and 130 of nsp12 over time (Figure 8K and L), suggesting that these mutations may play a crucial role in the dominance of these strains in the epidemic by enhancing the stability of nsp12.

**Figure 8.**
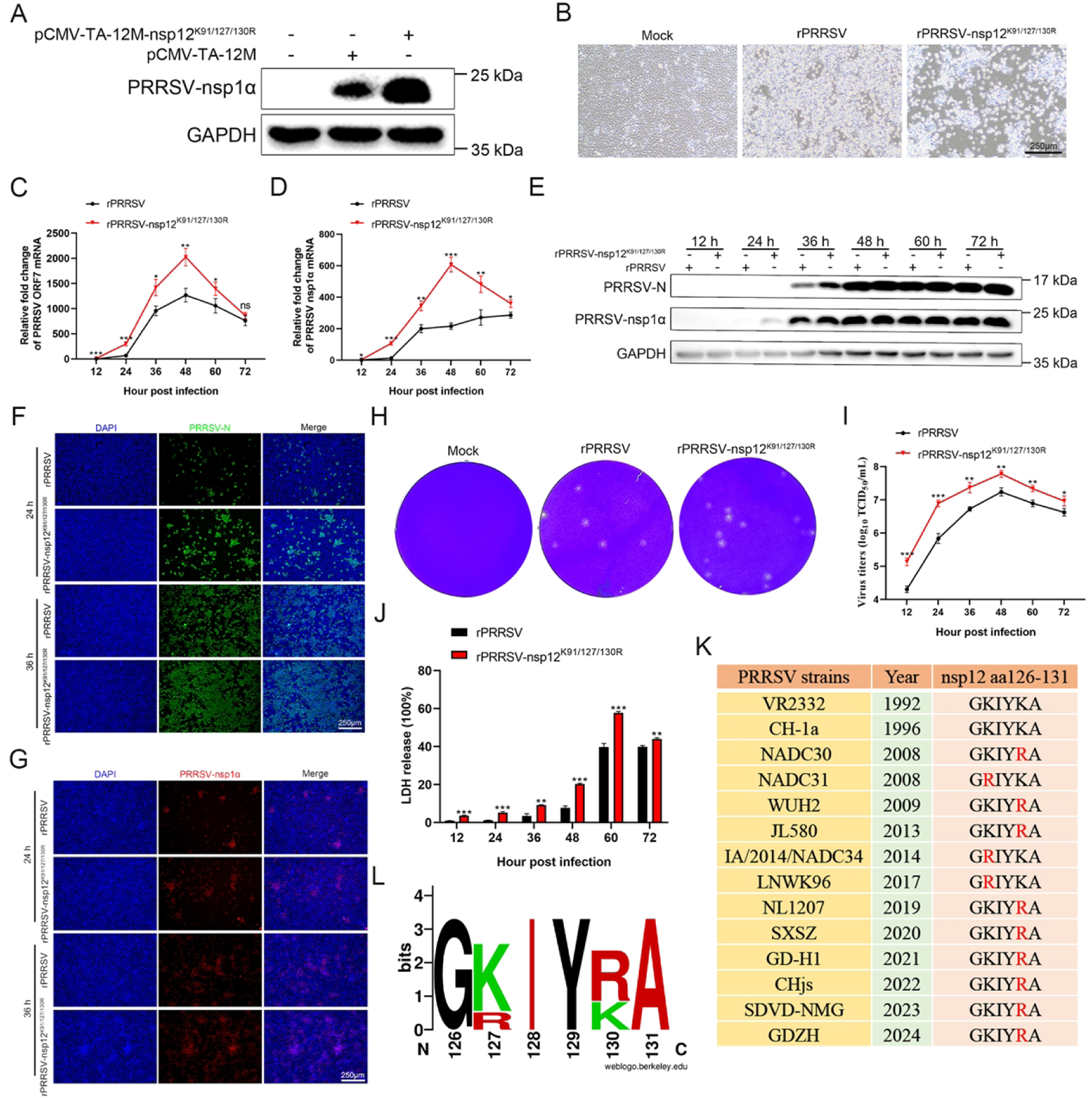
The rPRRSV-nsp12^K91/127/130R^ exhibits greater replication capacity compared to rPRRSV. (**A**) HEK293T cells in a 6-well plate were transfected with 2.5 μg cDNA clone plasmid using Lipofectamine 3,000. At 48 h, cells were harvested for nsp1α protein detection by western blotting. (**B**) Marc-145 cells in 12-well plates were infected with recombinant viruses at an MOI of 0.05 for 84 h. CPE was observed under a light microscope. (**C-E**) Marc-145 cells in 12-well plates were infected with recombinant viruses at an MOI of 0.05. The mRNA and protein levels of N and nsp1α were detected by RT-quantitative PCR (qPCR) and western blotting. (**F and G**) Marc-145 cells in 12-well plates were infected with recombinant viruses at an MOI of 0.05 for 24 and 36 h. The expression of N and nsp1α proteins was assessed by IFA. (**H**) Marc-145 cells in 12-well plates were infected with recombinant viruses at an MOI of 0.05 for 24 h, followed by a plaque assay. (**I and J**) Marc-145 cells in 12-well plates were infected with recombinant viruses at an MOI of 0.05. Culture supernatants were collected at 12, 24, 36, 48, 60 and 72 h, and viral titers (**I**) as well as LDH release (**J**) were measured. (**K**) Alignment of the amino acid sequences of positions 126-131 within nsp12 from different PRRSV strains. (**L**) The frequency of amino acid changes at positions 126-131 in PRRSV nsp12 (Figure 8K) was analyzed using the Weblogo3 online tool. Mean ± SEM of three independent experiments. *: *p* < 0.05; **: *p* < 0.01; ***: *p* < 0.001; ns: not significant (Student’s *t*-test).

### The rPRRSV-nsp12^R89K^ strain exhibits significantly lower replication capacity compared to rPRRSV

Notably, sequence alignment revealed that the TA-12 strain harbors a K89R substitution in nsp12 compared to the prototype VR2332 strain (Figure S4A). Given our previous demonstration that lysine at position 89 of nsp12 is a critical ubiquitination site, we hypothesized that reversion of this mutation (R89K) would restore ubiquitination-dependent degradation and attenuate viral replication. To test this, we generated a recombinant virus (rPRRSV-nsp12^R89K^) based on the TA-12 backbone (Figure S5A). The successful rescue of rPRRSV-nsp12^R89K^ was confirmed by CPE (Figure S5B), western blotting (Figure S5C), IFA (Figure S5D and E), RT-PCR (Figure S5F), and Sanger sequencing (Figure S5G). Further studies revealed that rPRRSV- nsp12^R89K^ induced weaker CPE compared to rPRRSV (Figure 9A). Correspondingly, the mRNA (Figure 9B and C) and protein (Figure 9D) levels, as well as the fluorescence intensity (Figure 9E and F) of N and nsp1α in rPRRSV-nsp12^R89K^, were significantly lower than in rPRRSV. Although both strains formed similar-sized plaques (Figure 9G), the viral titer of rPRRSV-nsp12^R89K^ was significantly lower (Figure 9H). Moreover, infection with rPRRSV-nsp12^R89K^ resulted in reduced LDH release compared to rPRRSV (Figure 9I). Strikingly, longitudinal sequence analysis indicated that lysine at position 89 of nsp12 was replaced by arginine over time (Figure 9J and K). These findings collectively demonstrate that the K89R substitution represents an evolutionary adaptation that *enhances viral fitness by evading host ubiquitin-mediated defense mechanisms*.

**Figure 9.**
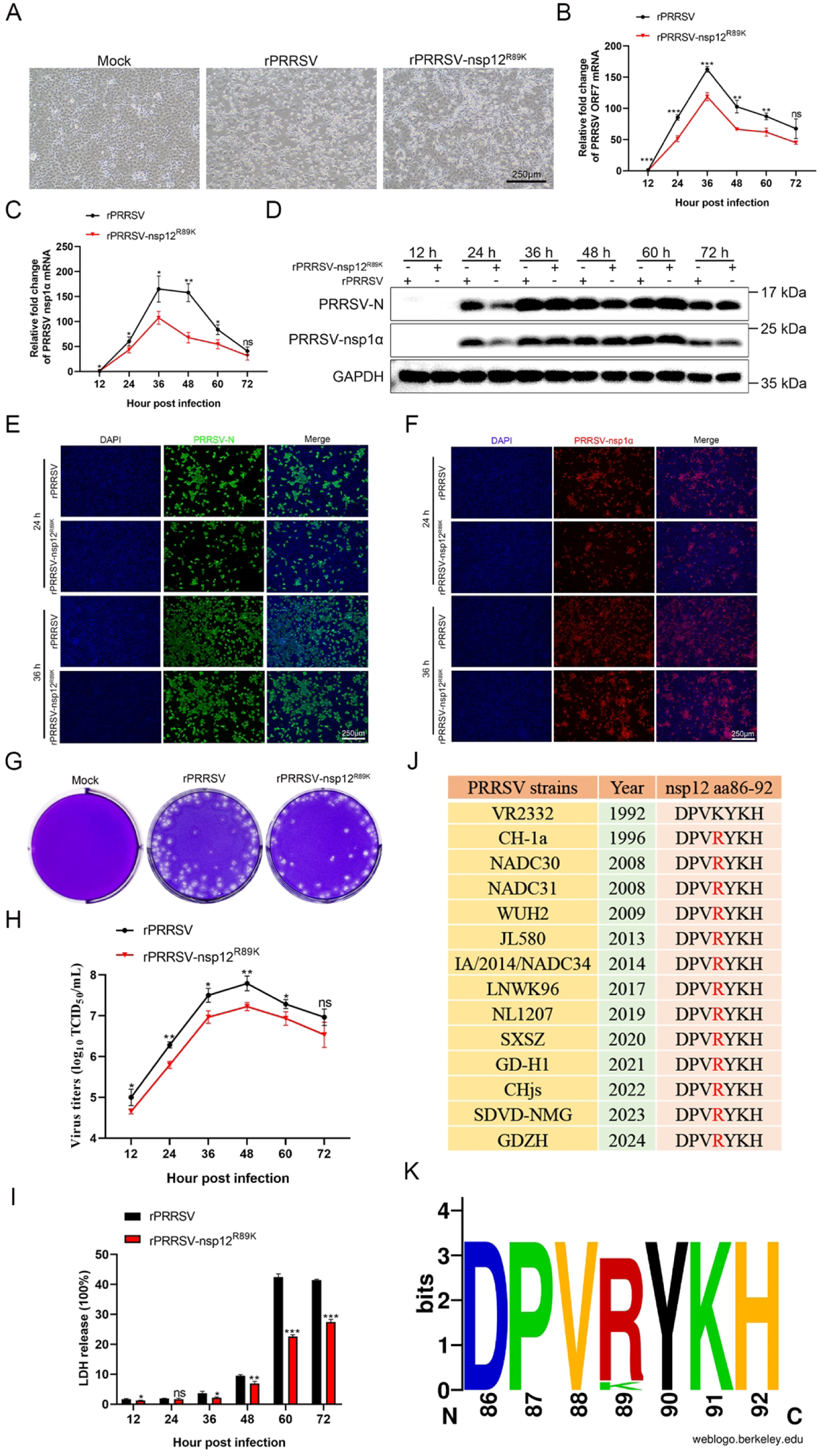
The rPRRSV-nsp12^R89K^ strain exhibits significantly lower replication capacity compared to rPRRSV. (**A**) Marc-145 cells in 12-well plates were infected with recombinant viruses at an MOI of 0.1 for 78 h. CPE was observed under a light microscope. (**B-D**) Marc-145 cells in 12-well plates were infected with recombinant viruses at an MOI of 0.1. The mRNA and protein levels of N and nsp1α were detected by qPCR and western blotting. (**E and F**) Marc-145 cells in 12-well plates were infected with recombinant viruses at an MOI of 0.1 for 24 and 36 h. The expression of N and nsp1α proteins was assessed by IFA. (**G**) Marc-145 cells in 12-well plates were infected with recombinant viruses at an MOI of 0.1 for 24 h, followed by a plaque assay. (**H and I**) Marc-145 cells in 12-well plates were infected with recombinant viruses at an MOI of 0.1. Culture supernatants were collected at 12, 24, 36, 48, 60 and 72 h, and viral titers (**H**) as well as LDH release (**I**) were measured. (**J**) Alignment of the amino acid sequences of positions 86-92 within nsp12 from different PRRSV strains. (**K**) The frequency of amino acid changes at positions 86-92 in PRRSV nsp12 (Figure 9J) was analyzed using the Weblogo3 online tool. Mean ± SEM of three independent experiments. *: *p* < 0.05; **: *p* < 0.01; ***: *p* < 0.001; ns: not significant (Student’s *t*- test).

### The nsp12 interactome is primarily associated with ubiquitination-related modifications

To elucidate the interaction network and regulatory mechanisms between nsp12 and host proteins during PRRSV infection, we performed liquid chromatography and tandem mass spectrometry (LC-MS/MS) using nsp12-mCherry as bait (Figure 10A). The Co-IP experiments were conducted to pull down the nsp12-mCherry complex and its interacting proteins from cell lysates using an anti-mCherry antibody. The results showed that the anti-mCherry antibody successfully pulled down the nsp12-mCherry complex, whereas the control anti-IgG antibody failed to do so (Figure 10B). The immunoprecipitated proteins were subsequently analyzed by mass spectrometry, which identified 176 proteins specifically interacting with nsp12-mCherry (Figure 10C, Files S1 and S2). To further explore the biological functions of these interacting proteins, Gene Ontology (GO) analysis was performed, revealing significant enrichment in genes related to ubiquitination modification (Figure 10D-F). This finding was further corroborated by KEGG pathway analysis, which demonstrated that nsp12-interacting proteins were predominantly associated with ubiquitination pathways (Figure 10G), suggesting a potential role for ubiquitin-mediated regulation in nsp12-host protein interactions.

**Figure 10.**
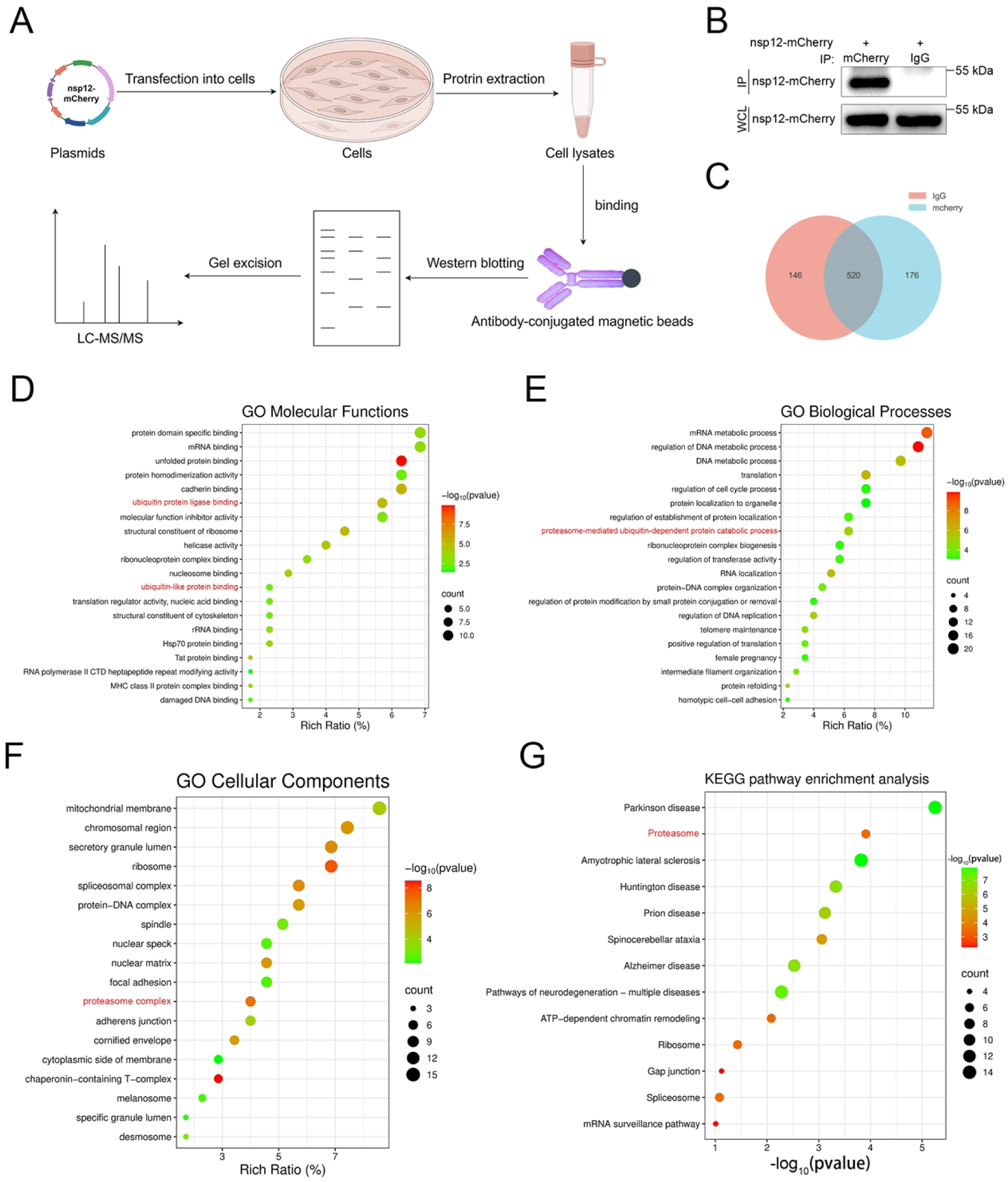
The nsp12 interactome is primarily associated with ubiquitination- related modifications. (**A**) Schematic diagram showing the IP-MS procedure for mapping the nsp12-host protein interactomes. (**B**) HEK293T cells were transfected with plasmids encoding nsp12-mCherry, followed by Co-IP using either anti-mCherry or control anti-IgG conjugated beads. The immunoprecipitated complexes were analyzed by both western blotting and LC-MS/MS. (**C**) Identification of proteins specifically interacting with nsp12 by a Venn diagram in Venny. (**D-F**) GO function analysis. (**G**) KEGG pathway enrichment analysis.

## Discussion

Ubiquitination plays a crucial role in regulating host innate immune responses and mediating the selective degradation of viral components to inhibit viral proliferation. However, numerous viruses have evolved mechanisms to manipulate the ubiquitination system, suppressing host defense responses to promote their own replication [32,33]. In this study, we uncovered a novel molecular mechanism underlying the antagonistic interplay between the host and the virus. We demonstrate that PRRSV nsp12 undergoes host-mediated ubiquitination and subsequent degradation via the ubiquitin-proteasome pathway and selective autophagy. However, nsp12 escapes host-mediated degradation through adaptive mutations at lysine residues, effectively circumventing host antiviral defenses. These results demonstrate that host ubiquitination enhances antiviral capacity, while the virus maintains its survival advantage through mutations, revealing a dynamic evolutionary arms race between the host and the virus.

As a key post-translational modification, ubiquitination regulates protein stability, localization, and function, playing critical roles in physiological and pathological processes [34,35]. As a central modulator of host-virus interactions, it impacts virus replication, immune evasion, and pathogenesis [36–38]. This modification not only enhances antiviral responses by activating host immune signaling pathways, such as the RIG-I-like receptor (RLR), the cyclic GMP-AMP synthase (cGAS)-stimulator of interferon genes (STING), and the nuclear factor kappa-B (NF-κB) pathways [39–43], but also directly targets viral proteins for degradation, thereby restricting viral replication [44–46]. However, many viruses have evolved mechanisms to exploit the host ubiquitination system, either by manipulating host signaling pathways or promoting the degradation of antiviral proteins, facilitating their replication and persistence [47–51]. Understanding the molecular mechanisms behind viral manipulation of the ubiquitin-proteasome system can lead to the identification of novel antiviral drug targets and the development of new strategies to enhance the host’s antiviral responses.

PRRSV, characterized by high virulence and continuous emergence of genetically divergent variants, represents a persistent challenge to the global swine industry [52]. Understanding the mechanisms underlying PRRSV pathogenesis is essential for developing efficacious vaccines and antiviral strategies. Post-translational modifications, particularly ubiquitination, play a critical role in regulating viral replication, immune evasion, and host antiviral responses, making them promising therapeutic targets [53]. Recent studies have highlighted the importance of ubiquitination in PRRSV infection. For example, USP1 deubiquitinates PRRSV nsp1β at K48-linked chains, stabilizing the protein and enhancing viral replication [54]. Additionally, NLRP12 promotes K48-linked ubiquitination of GP2a at K128, leading to its degradation via the lysosomal pathway mediated by MARCH8-NDP52 [55]. While the nsps of PRRSV are known to play a pivotal role in viral replication and virulence [14,56,57], the role of host-mediated ubiquitination in modulating the stability of nsps and their influence on viral replication remains poorly defined. In this study, we observed that nsp12 exhibits instability and is susceptible to degradation through the host’s proteasome pathway (Figure 1). Ubiquitination modification exhibits diverse functions, among which K48- and K63-linked chains are the most abundant linkage types. Here, we found that nsp12 underwent both K48- and K63-linked polyubiquitination, suggesting its potential degradation through these distinct ubiquitin-dependent pathways (Figure 1). Previous studies have shown that PSMB1- mediated degradation of nsp12 primarily depends on its ubiquitination at lysine residue 130 [58]. However, our research revealed that host proteins could catalyze ubiquitination of nsp12 at multiple sites. Further analysis indicated that lysine residues 89, 91, 127, and 130 were the main ubiquitination sites for nsp12 (Figure 2). These results suggest that, in addition to PSMB1, other host proteins may also regulate the stability of nsp12 by mediating its ubiquitination, thereby affecting the viral replication process. Previous studies reported that RNF114 promotes nsp12 ubiquitination, leading to its proteasomal degradation, though the underlying mechanism remained unclear [30]. In this study, we demonstrated that RNF114-dependent degradation of nsp12 required its E3 ligase activity. Crucially, mutation of the catalytic site in RNF114 significantly impaired its ability to suppress PRRSV replication, consistent with findings that RNF114’s E3 activity is essential for inhibiting CSFV replication [31].

Furthermore, we identified lysine residues K127 and K130 of nsp12 as the primary ubiquitination sites targeted by RNF114 (Figure 3).

Ubiquitination serves as a central regulatory nexus, coordinating both proteasomal degradation and selective autophagy pathways [29]. In addition to its role in the clearance and recycling of cellular components, autophagy also plays a critical role in host defense by targeting various invading pathogens, such as bacteria and viruses [59–61]. Previous studies have shown that autophagy receptors can mediate the selective autophagic clearance of viral proteins through ubiquitin-dependent mechanisms. This process has been evolutionarily conserved across diverse viruses, as evidenced by studies on pseudorabies virus (PRV), hepatitis B virus (HBV), porcine epidemic diarrhea virus (PEDV), and Seneca Valley virus (SVV) [62–65]. Similarly, studies have reported that PSMB4 inhibits PRRSV replication by degrading nsp1α through selective autophagy [66]. In this study, we demonstrated that exogenous elevation of ubiquitin levels promoted the degradation of nsp12, a process that was rescued by the autophagy inhibitor 3-MA. However, when all lysine residues within nsp12 were substituted with arginine, 3-MA treatment no longer affected its expression, establishing lysine residues as indispensable determinants for host-mediated autophagic degradation (Figure 4). In contrast to non-selective autophagy, selective autophagy relies on specific autophagy receptors (such as NBR1, SQSTM1/p62 and others), which recognize ubiquitin-tagged substrates and bind them to the autophagosome marker protein LC3, facilitating targeted delivery to autolysosomes for degradation [67]. Notably, we found that a multi-receptor recognition system involving NBR1, OPTN, P62, NDP52 and TOLLIP interacted with nsp12, suggesting its potential targeting for selective autophagic degradation (Figure 5). Further studies revealed that nsp12 forms complexes with autophagy receptors and LC3, and knockdown of NBR1, SQSTM1, or NDP52 significantly inhibited host-mediated degradation of nsp12 (Figure 6). This finding is consistent with previous reports that PSMB1 recruits NBR1 to mediate the autophagic degradation of nsp12, further supporting the critical role of autophagy receptors in nsp12 degradation [58].

RNA viruses, characterized by their high mutation rates and recombination capacity, present formidable obstacles for epidemiological containment [68,69]. Furthermore, to adapt to environmental pressures and evade host defense mechanisms, viral proteins often undergo adaptive mutations at specific amino acid sites, thereby enhancing their survival and replication capabilities [70–72]. To investigate whether nsp12 of PRRSV escapes host degradation mechanisms through adaptive mutations in its amino acid sequence, we conducted an evolutionary analysis of its amino acid sequence. Our analysis indicated that the lysine residue within nsp12 showed a tendency to arginine during evolution (Figure 7). To further investigate whether nsp12 ubiquitination affects viral replication, a recombinant PRRSV (rPRRSV-nsp12^K91/127/130R^) strain was constructed. Results showed this strain exhibited significantly enhanced replication capacity *in vitro*. Conversely, the rPRRSV-nsp12^R89K^ strain showed markedly reduced infectivity *in vitro* (Figures 8 and 9). In fact, while the VR2332 strain initially contains lysine at nsp12 position 89, evolutionary tracking revealed progressive replacement by arginine (Figure 9). Concurrently, lysine residues at positions 91 and 130 also displayed an evolutionary trend toward arginine substitution (Figure 8). These findings suggest that PRRSV escapes host-mediated ubiquitination-dependent degradation by mutating lysine to arginine, thereby enhancing viral fitness. Similar immune evasion strategies have been reported in other viruses. For instance, influenza virus (IAV) escapes MARCH8-mediated host restriction by mutating lysine at position 78 of the M2 protein [73]. In addition, mass spectrometry analysis demonstrated that nsp12-interacting proteins were significantly enriched in ubiquitination-related modifications (Figure 10). However, the functional relationships between these proteins and nsp12, their regulatory roles, and the underlying mechanisms require further investigation.

In summary, our study reveals a dynamic antagonistic interplay between the host and the virus. The host degrades nsp12 through the ubiquitin-proteasome pathway and selective autophagy to inhibit viral replication, while the virus escapes host-mediated degradation via adaptive mutations within nsp12 (Figure 11). These findings not only deepen our understanding of PRRSV pathogenesis but also provide a theoretical foundation for developing novel anti-PRRSV strategies targeting the regulation of ubiquitination and autophagy.

**Figure 11.**
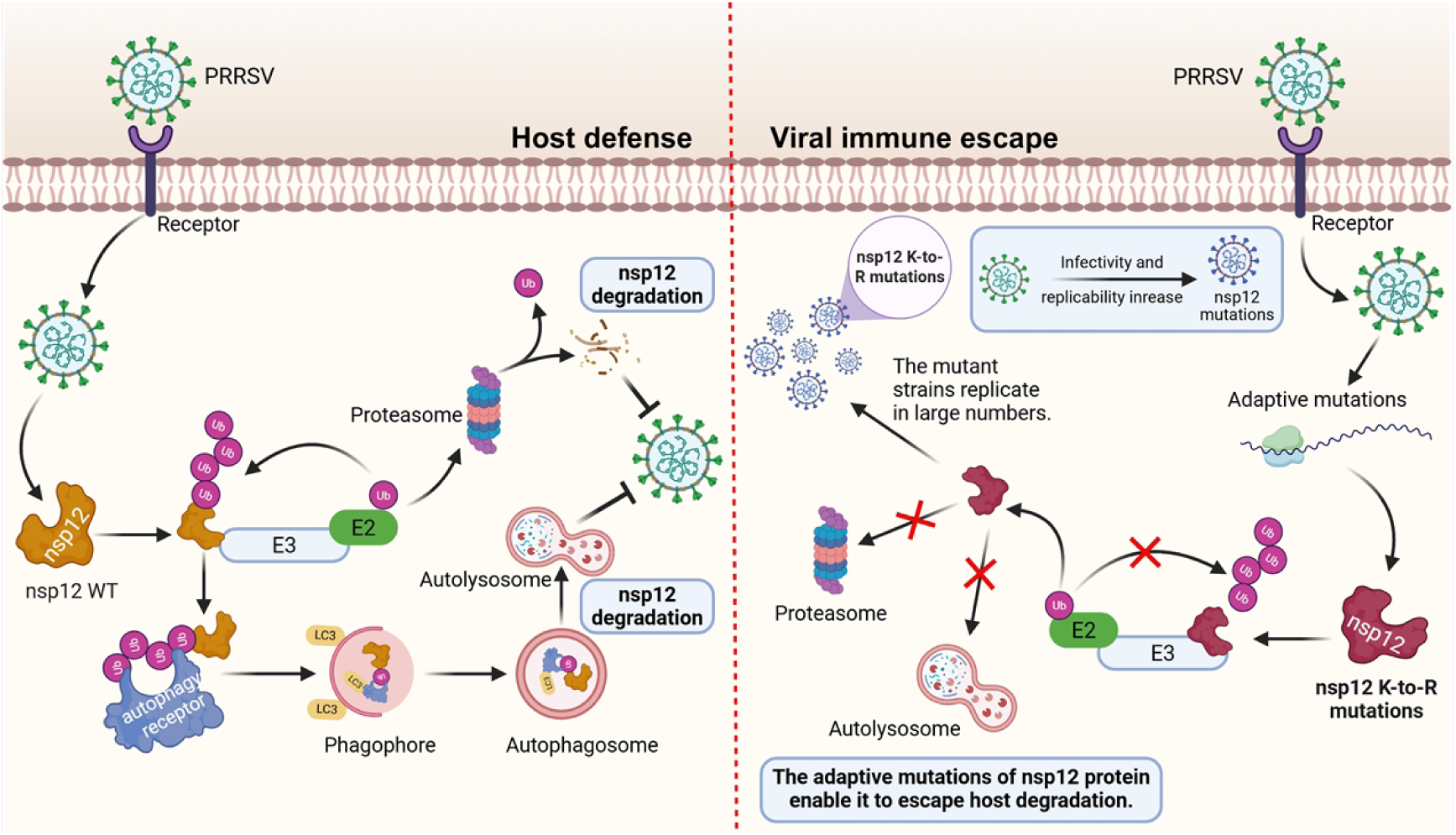
A model demonstrating how adaptive lysine mutations within PRRSV nsp12 evade host proteasomal and selective autophagic degradation. Upon PRRSV infection, cellular E3 ubiquitin ligases specifically interact with and mediate the ubiquitination of viral nsp12 protein, which is subsequently degraded through the proteasomal pathway and ubiquitin-dependent selective autophagy, ultimately suppressing viral replication. However, evolutionary pressure drives the accumulation of lysine-to-arginine substitutions at critical residues within nsp12. These mutations abrogate ubiquitination-dependent degradation of nsp12, resulting in protein stabilization and increased viral replication efficiency.

## Materials and methods Cells and viruses

African green monkey kidney cells (Marc-145) and HEK293T cells were cultured in Dulbecco’s modified Eagle’s medium (DMEM; Gibco, 8122784) supplemented with 10% fetal bovine serum (FBS; ExCell Bio, FSP500). All cells were cultured at 37 ℃ with 5% CO_2_. PRRSV strains CH-1a (AY032626.1), TA12 (HQ416720.1), JXA1(FJ548855.1), SD16 (JX087437.1), and WUH3 (HM853673.2) were preserved in our laboratory. The recombinant strain rHP-PRRSV TA-12 (rPRRSV) was generated using a method similar to that described previously, with minor modifications [74]. PRRSV was propagated and titrated on Marc-145 cells.

## Antibodies and reagents

The mouse monoclonal antibody specific to PRRSV N (JN0401, 1:3000) was purchased from MEDIAN Diagnostics (Korea). Anti-PRRSV nsp1α antibody (GTX133695, 1:3000) was purchased from GeneTex. Anti-mCherry-tag (26765-1-AP, 1:2000), anti-GAPDH (60004-1-Ig, 1:5000), anti-ubiquitin (Ub) (10201-2-AP, 1:1000), anti-NBR1(16004-1-AP, 1:500), anti-OPTN (10837-1-AP, 1:2000), anti-TOLLIP (11315-1-AP, 1:3000), anti-SQSTM1 (18420-1-AP, 1:2000), anti-NDP52 (12229-1-AP, 1:1000), anti-GFP (66002-1-Ig, 1:2000), HRP-conjugated anti-mouse (SA00001- 1, 1:10000) or -rabbit (SA00001-2, 1:10000) and Alexa Fluor 647-conjugated goat anti-rabbit (RGAR005, 1:150) antibodies were purchased from Proteintech Group. Anti-HA-tag (3724, 1:3000), Alexa Fluor 488-conjugated goat anti-rabbit (4412, 1:1000) or -mouse (4408, 1:1000) and Alexa Fluor 555-conjugated goat anti-rabbit (4413, 1:1000) or -mouse (4409, 1:1000) antibodies were purchased from Cell Signaling Technology. Chemical reagents MG132 (HY-13259), 3-methyladenine (3-MA; HY-19312), and cycloheximide (CHX; HY-12320) were purchased from MedChemExpress. 4ʹ,6- diamidino-2-phenylindole, dihydrochloride (DAPI; C0065, 1:1000) was purchased from Solarbio. Protein A/G Magnetic Beads (B23202) were purchased from Selleck Chemicals. Lipofectamine 3000 (Invitrogen, L3000015) was purchased from Thermo Fisher Scientific.

## Plasmid construction and cell transfection

The nsp12 from the PRRSV VR2332 genome was subcloned into pmCherry-N1 vector to generate the nsp12-mCherry fusion expression plasmid. Nsp12 mutants were generated from the nsp12-mCherry by site-directed mutagenesis or chemical synthesis. HA-tagged ubiquitin mutants (K6, K11, K27, K29, K33, K48, K63, K48R and K63R) were generated from HA-Ub through point mutations. Myc-tagged RNF114 was constructed in the pcDNA3.1-Myc vector. HA-tagged NBR1, OPTN, SQSTM1, NDP52 and TOLLIP were cloned into the pCAGGS-HA vector. LC3 was amplified from HEK293T cells cDNA and then inserted into the pCDNA3.1-GFP vector to generate LC3-GFP. All constructed plasmids were verified by Sanger sequencing. For cell transfection, cells grown in culture plates were transfected at 80% confluence using either polyethylenimine (PEI; Servicebio, G1802) or Lipofectamine 3000, at a ratio of 1 µg plasmid to 3 µL transfection reagent.

## MG132, CHX and 3-MA treatment

For MG132 and 3-MA treatments, HEK293T cells were transfected with plasmids for 12 h before the addition of inhibitors. After a 12 h incubation, cells were harvested and lysates were analyzed by immunoblotting. For the protein half-life assay, HEK293T cells were transfected with plasmids for 24 h, and then treated with the protein synthesis inhibitor CHX (50 μg/mL). Cells were collected at the indicated time points, and target protein levels were analyzed by western blotting.

## Western blotting and Co-IP assay

For the western blotting assay, cells were lysed on ice for 30 min using IP cell lysis buffer (Beyotime, P0013J) supplemented with 1 mM PMSF protease inhibitor (Solarbio, P0100). Proteins were resolved by sodium dodecyl sulfate polyacrylamide gel electrophoresis (SDS-PAGE) and transferred electrophoretically onto polyvinylidene difluoride (PVDF) membranes (Merck Millipore, ISEQ00010). Membranes were blocked with 5% skim milk (Beyotime, P0216) for 2 h at room temperature, followed by overnight at 4°C with primary antibody and 1 h incubation at room temperature with HRP-conjugated anti-rabbit/mouse secondary antibodies. Protein bands were visualized using a Chemiluminescence Detection Kit (MeilunBio, MA0186-3).

For Co-IP assay, magnetic beads were conjugated to specified antibodies via 30 min incubation at room temperature. Cell lysate supernatants were incubated with antibody-conjugated beads for 2 h at room temperature or overnight at 4 ° C. Bead-bound complexes were washed three times, transferred to fresh tubes, and washed three additional times. Proteins were eluted by boiling in protein loading buffer for western blotting analysis.

## Indirect immunofluorescence assay

After washing with PBS, cells were fixed with 4% paraformaldehyde (Beyotime, P0099) for 15 min at room temperature, permeabilized with 0.5% Triton X-100 (Beyotime, P0096) for 5 min, and blocked with 1% bovine serum albumin (BSA; Beyotime, ST2249) for 1 h at room temperature. Cells were then incubated overnight with primary antibodies. After washing to remove unbound primary antibodies, cells were incubated with appropriate fluorescently labeled secondary antibodies for 1 h at room temperature. Following secondary antibody removal and washing, nuclei were counterstained with DAPI for 5 min. Fluorescence images were acquired using either an inverted fluorescence microscope (Nikon ECLIPSE Ti2) or a confocal laser scanning microscope (Leica, SP8).

## Yeast two-hybrid (Y2H) screening

For yeast two-hybrid (Y2H) screening, PRRSV nsp12 was cloned into the pGADT7 vector to generate the prey plasmid pGADT7-nsp12. NBR1, OPTN, SQSTM1, NDP52, TOLLIP, and LC3 were cloned into the pGBKT7 vector to construct the corresponding bait plasmids. The bait plasmids were cotransformed with pGADT7-nsp12 into yeast strain Y2HGold. Following transformation, positive colonies were selected and streaked onto SD/-Leu/-Trp/-His/-Ade/X-α-Gal/AbA (QDO/X/A) plates for further screening. In the Y2H assay, the combination of pGBKT7-53 and pGADT7-T served as the positive control. Conversely, the pairing of pGBKT7-Lam and pGADT7-T was used as the negative control.

## RNA interference (RNAi) assay

Small interfering RNAs (siRNAs) targeting human NBR1, OPTN, SQSTM1, NDP52 and TOLLIP, along with scramble siRNA controls, were synthesized by IGE biotech (Guangzhou, China). HEK293T cells were transfected with siRNAs for 12 h using Lipofectamine 3000, followed by transfection with other specified plasmids using PEI for 24 h. The siRNA sequences are listed in Supplementary Table S3.

## Phylogenetic and sequence analyses of the PRRSV nsp12

For phylogenetic analysis, a neighbor-joining (NJ) method in MEGA software was used to construct phylogenetic trees with 1,000 bootstrap replicates. Tree visualization, manipulation, and annotation were performed using the Interactive Tree of Life platform (iTOL; https://itol.embl.de). For sequence analysis, nucleotide and amino acid homology of the nsp12 was analyzed with the Clustal W method in MegAlign program of the DNASTAR software package (version 7.0; DNASTAR, Madison, WI). Sequence alignment and site-specific variation analysis of the nsp12 amino acid sequences were conducted using BioEdit software (version 7.2.6.1). The frequency of lysine mutations in PRRSV nsp12 was counted using WebLogo3 (http://weblogo.berkeley.edu/).

## RNA extraction and quantitative real-time PCR

Total RNA was extracted from Marc-145 cells using VeZol Reagent (Vazyme, R411) and reverse-transcribed into complementary DNA (cDNA) using the HiScript II Q RT SuperMix for qPCR Kit (Vazyme, R223). Quantitative real-time PCR was performed with the ChamQ Blue Universal SYBR qPCR Master Mix (Vazyme, Q312) on a QuantStudio ™ 3 instrument (Applied Biosystems, USA). Gene expression changes were calculated via the 2^- ΔΔCt^ method using *GAPDH* as the endogenous control. All primer sequences are listed in Supplementary Table S4.

## Plaque Assay

Marc-145 cells were cultured in 6-well plates until reaching 100% confluency. The cells were infected with a 10-fold serial dilution of PRRSV for 1 h, with gentle mixing every 15 min. After infection, the culture medium was removed, and cells were washed twice with PBS. Subsequently, cells were overlaid with DMEM containing 1% low-melting agarose (Sangon Biotech, A600015) and 2% FBS. After agarose solidification at room temperature, plates were inverted and incubated at 37°C with 5% CO₂. At 72 h after infection, the agarose layer was carefully removed and the plaques were visualized after staining with crystal violet solution.

## Cell viability assay

Marc-145 cells were cultured in a 12-well plate and infected with PRRSV. Cell culture supernatants were collected at 12, 24, 36, 48, 60, and 72 h. LDH release was measured using the CytoTox 96® Non-Radioactive Cytotoxicity Assay (Promega, G1780) according to the manufacturer’s instructions.

## Construction and rescue of recombinant viruses

The highly pathogenic rTA-12 strain was used as the backbone to construct recombinant viruses carrying the K91/127/130R or R89K mutations in nsp12, following a procedure similar to that previously described [74]. Briefly, HEK293T cells were cultured in six-well plates to 80% confluence, then transfected with 2.5 μg of infectious full-length cDNA clones using Lipofectamine 3000. At 48 h post-transfection, culture supernatants were collected as passage zero (P0) virus and used to infect Marc-145 cells. When CPE became evident, both cells and supernatants were harvested as P1 virus and subjected to freeze-thaw cycles for propagation.

## Functional enrichment analysis of interacting proteins

Functional annotation of nsp12-interacting proteins was performed through GO term enrichment using the Metascape website (https://metascape.org/gp/index.html), while KEGG pathway analysis was conducted using DAVID Bioinformatics Resources (http://david.ncifcrf.gov). The enriched functional terms were then analyzed and visualized using the bioinformatics online platform (https://www.bioinformatics.com.cn/).

## Statistical analysis

All data were analyzed using GraphPad Prism 8 software. The differences between two groups were assessed using Student’s t-test, while one-way or two-way ANOVA was used for multiple comparisons. A *P*-value < 0.05 was considered statistically significant (\**P* < 0.05, \*\**P* < 0.01, \*\*\**P* < 0.001), and "ns" indicates no significant difference.

## Acknowledgements

We are very grateful to Prof. Yanhua Li from Yangzhou University, China for providing the infectious cDNA clone of TA-12 (pCMV-TA-12 M).

## Author Contributions

**Yongjie Chen:** Conceptualization, Methodology, Data curation, Validation, Visualization, Writing - original draft. **Zhan He:** Methodology, Investigation. **Zishen Chen:** Methodology, Investigation. **Ling Huang:** Methodology, Investigation. **Haotong Lu:** Investigation, Validation. **Siyong Zeng:** Investigation, Validation. **Baoying Huang:** Investigation, Validation. **Chunhe Guo:** Conceptualization, Funding acquisition, Supervision, Writing - review & editing.

## Data availability statement

All data supporting the conclusions are included in this article and its supplementary materials. Other source data provided by our findings are also available from the corresponding author upon reasonable request.

